# Deletion of *TRPC6*, an autism risk gene, induces hyperexcitability in cortical neurons derived from human pluripotent stem cells

**DOI:** 10.1101/2022.11.14.516407

**Authors:** Kyung Chul Shin, Gowher Ali, Houda Yasmine Ali Moussa, Vijay Gupta, Alberto de la Fuente, Hyung-Goo Kim, Lawrence W Stanton, Yongsoo Park

**Author notes:** Corresponding authors; Dr. Yongsoo Park, Neurological Disorders Research Center, Qatar Biomedical Research Institute (QBRI), Hamad Bin Khalifa University (HBKU), Qatar Foundation, Doha, Qatar, Dr. Lawrence W Stanton, Neurological Disorders Research Center, Qatar Biomedical Research Institute (QBRI), Hamad Bin Khalifa University (HBKU), Qatar Foundation, Doha, Qatar. These authors contributed equally to this work (Kyung Chul Shin, Gowher Ali). **Competing Interests:** The authors declare no competing interests. **Author Contributions** Conceptualization: K.C.S., L.W.S., Y.P.; methodology and investigation: K.C.S., G.A., V.G., A.F., H.Y.A.M.; visualization: K.C.S., G.A., V.G., A.F.,Y.P.; project administration and funding acquisition: L.W.S., Y.P.; supervision: H.G.K., L.W.S., Y.P.; writing - original draft: K.C.S., L.W.S., Y.P.

## Abstract

Autism spectrum disorder (ASD) is a complex and heterogeneous neurodevelopmental disorder linked to numerous rare, inherited and arising *de novo* genetic variants. ASD often co-occurs with attention-deficit hyperactivity disorder and epilepsy, which are associated with hyperexcitability of neurons. However, the physiological and molecular mechanisms underlying hyperexcitability in ASD remain poorly understood. Transient receptor potential canonical-6 (TRPC6) is a Ca^2+^-permeable cation channel that regulates store-operated calcium entry (SOCE) and is a candidate risk gene for ASD. Using human pluripotent stem cell (hPSC)-derived cortical neurons, single cell calcium imaging, and electrophysiological recording, we show that TRPC6 knockout (KO) reduces SOCE signaling and leads to hyperexcitability of neurons by increasing action potential frequency and network burst frequency. Our data provide evidence that reduction of SOCE by TRPC6 KO results in neuronal hyperexcitability, which we hypothesize is an important contributor to the cellular pathophysiology underlying hyperactivity in some ASD.

## Introduction

Autism spectrum disorder (ASD) is a complex neurodevelopmental disorder, characterized by stereotyped repetitive behaviors and communication deficits^1^. An increasing numbers of genetic variants implicated in ASD have been reported, suggesting a high degree of locus heterogeneity and a contribution from rare and *de novo* variants^2^. Comorbidity is common in ASD, including attention-deficit hyperactivity disorder (ADHD) and epilepsy, which are associated with hyperexcitability of neurons^3^. ASD is phenotypically and etiologically so heterogeneous that it is challenging to determine a contributing role of various variants on ASD etiology and to uncover the underlying cellular and molecular pathophysiology. The functional study of genetic variants associated with ASD is critical for the elucidation of ASD pathophysiology, thereby moving from gene discovery to understanding the biological influences of genetic variants for the development of ASD therapeutics^4^.

Risk variants associated with ASD converge on common cellular signaling and molecular pathways in neurons^5^. Intracellular calcium signaling is dysregulated in ASD and risk variants of ASD may cause deleterious effects on calcium signaling of the endoplasmic reticulum (ER), a major calcium store^6^. However, it remains poorly understood how calcium dysregulation gives rise to the pathophysiology of ASD and the hyperexcitability phenotype of ASD.

Calcium ions (Ca^2+^) are second messengers that control diverse biological processes, including both short-term response on neurotransmission and long-term effects on gene expression and neuronal differentiation^7, 8, 9^. Store-operated Ca^2+^ entry (SOCE) is the process by which the emptying of ER calcium stores causes influx of calcium^10^ and maintains calcium homeostasis through the connection of the ER/plasma membrane^11^. SOCE regulates neuronal signaling required for the maintenance of spines, neuronal excitability, and gene transcription^12^. SOCE is mainly mediated by ORAI, a calcium channel in the plasma membrane, and STIM (stromal interaction molecules), an ER calcium sensor^13, 14, 15, 16^. Dysregulation of SOCE is linked to neurological disorders such as Alzheimer’s disease, Huntington’s disease, and Parkinson’s disease^12, 17^.

Transient receptor potential canonical (TRPC) channels are a family of Ca^2+^-permeable cation channels that regulates SOCE by modulating STIM1 activity and the ternary complex of STIM1– ORAI1–TRPC^18^. TRPC6 is a candidate risk factor for ASD and implicated in ASD etiology: de novo missense and nonsense mutations in TRPC6 associated with ASD etiology have been reported^19, 20, 21^. Loss-of-function mutations in TRPC6 reduce calcium influx in human pluripotent stem cell (hPSC)-derived neurons^19^ and TRPC6 knockdown (KD) in *Drosophila* causes autism-like behavioral deficits and leads to a hyperactivity phenotype^21^. However, the pathophysiology underlying hyperactivity phenotype caused by TRPC6 KD in ASD is unclear.

A major impediment to ASD research is the lack of relevant animal and cellular models. Reprogramming somatic cells to a pluripotent state enables the development of neuronal models to study human diseases^22^. Patient-specific hPSC-derived neurons recapitulate the genomic, molecular and cellular attributes of developing native human neuronal subtypes with advantages over single time point studies. TRPC6 knockout (KO) hPSC lines were generated using CRISPR/Cas9 genome-editing techniques. We validated that hPSC-derived cortical neurons are functionally active. TRPC6 KO reduced SOCE calcium signaling and caused hyperexcitability of neurons by increasing network burst frequency and action potential frequency. Taken together, our data unveil the molecular and cellular pathophysiology underlying hyperactivity of ASD individuals. TRPC6 KO hPSC-derived cortical neurons reproduce an ASD hyperexcitability phenotype and thus provide a platform to model ASD neuropathology and pave the way for further studies to discover therapeutics for intervention of ASD.

## Results

### Generation of TRPC6 KO hPSC-derived cortical neurons

TRPC6 mutations in ASD individuals are genetic risk factors for ASD^19, 20, 21^ and TRPC6 KD causes the autism-like hyperactivity behavior in *Drosophila*^21^. To study the pathophysiology of ASD in a human neuron-based model, we generated TRPC6 KO hPSC lines by CRISPR-Cas9 gene editing (**Supplementary Fig. 1**). The guide RNA was designed to target the genomic sequence of TRPC6 exon 1, upstream of translation start site (**Supplementary Fig. 1a**). Two TRPC6 KO clones, termed C21 and C47, were established, where 65 bp and 49 bp were deleted, respectively including the region of the start codon (**Supplementary Fig. 1b-d**). The TRPC6 KO hPSCs showed normal morphology similarly with wild-type control hPSCs (**Supplementary Fig. 2a**) and expressed pluripotency markers including OCT4, NANOG, and SOX2 in monolayer cultured on Matrigel in mTesR1 medium (**Supplementary Fig. 2b**).

For neuronal differentiation, we utilized and modified previously described methods^23^ that creates functional cortical neurons with greater than 90% efficiency (**Fig. 1a,b**). We established the protocol for differentiation of hPSCs to mature human cortical neurons (**Fig. 1b**). Wild-type (CRTD5) hPSC lines were differentiated to neural progenitor cells (NPCs) (Week 0), and then NPCs further differentiated to cortical neurons up to 8 weeks (**Fig. 1b**). NPCs generated self-organizing rosette structures that expressed sex-determining region Y-related HMG box 2 (SOX2), a marker for neural stem cells and neural progenitors as well as telencephalic markers FOXG1 and OTX2, suggesting efficient neural conversion (**Supplementary Fig. 2c**).

**Figure 1.**
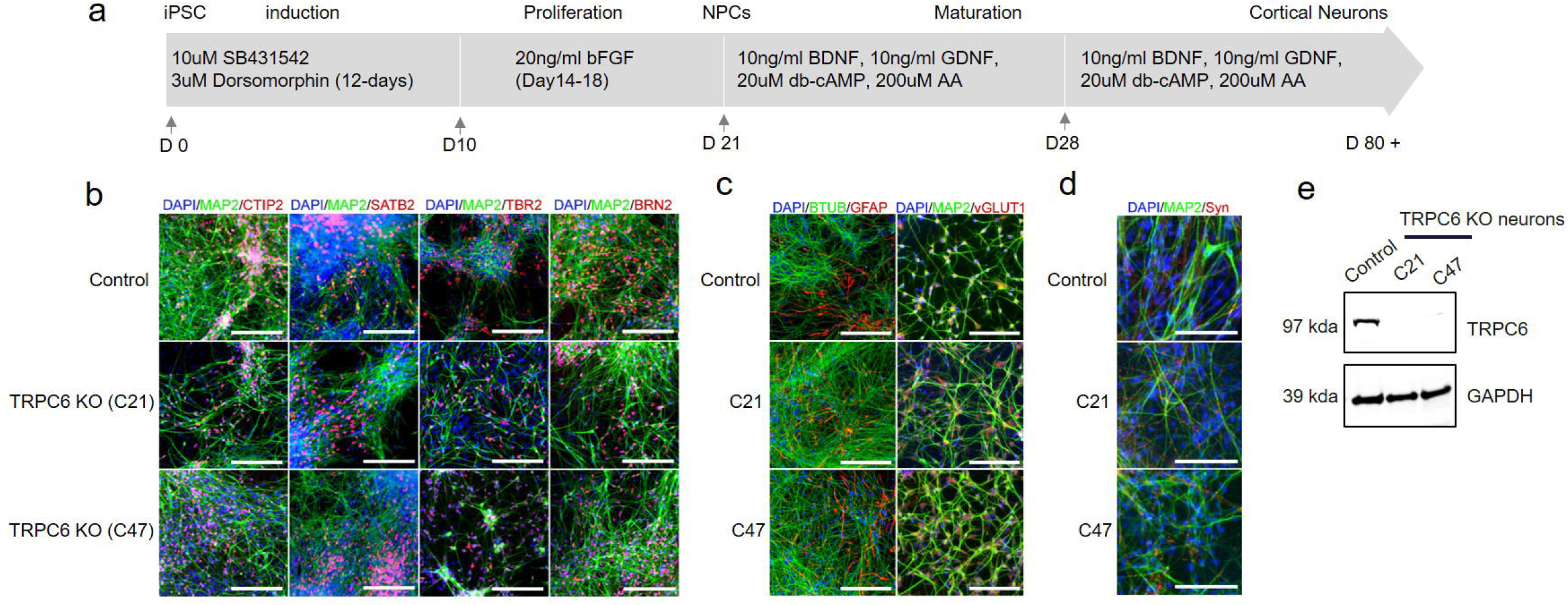
Characterization of human cortical neurons generated *in-vitro* from pluripotent stem cells. (**a**) Schematic of hPSC differentiation into NPCs and mature cortical neurons. The arrows indicate the time point of splitting the cells. (**b**) Immunostaining of control and TRPC6 KO cortical neurons (from two different hPSC clones, C21 and C47) differentiated for 8 weeks. Cortical neuron markers; CTIP2, SATB2, TBR2, and BRN2. MAP2, BTUB; pan-neuronal markers. (**c**) GFAP, a marker for glia. vGLUT1, a marker for glutamatergic excitatory synapses. (**d**) Synaptophysin (Syn), a synaptic vesicle protein. Nuclei are stained with DAPI. (**e**) Validation of TRPC6 protein expression in TRPC6 KO cortical neurons differentiated for 8 weeks. Scale, 100 μm.

Human cortical neurons after 8 weeks of differentiation were immunostained using antibodies against cortical neuron markers. Neurons expressed microtubule associated protein-2 (MAP2, a pan-neuronal marker) together with cortical upper layer markers such as CTIP2/BCL11B, special AT-rich sequence-binding protein 2 (SATB2), and POU domain, class 3, transcription factor 2 (POU3F2, also known as BRN2) as well as T-box brain protein 2 (TBR2, a cortical deep layer marker), showing the efficient differentiation of cortical neurons with characteristics of both upper and deep cortical layers (**Fig. 1b**). Given the low number of cells expressing glial fibrillary acidic protein (GFAP), a marker for astrocytes, with beta tubulin III (BTUB, a neuron marker) at 8 weeks of differentiation from NPCs, we estimated the efficiency of neuronal differentiation was >90% (**Fig. 1c**). Vesicular glutamate transporter 1 (vGLUT1, a marker for glutamatergic excitatory synapses) and synaptophysin (Syn, a synaptic vesicle protein) were colocalized with MAP2 (**Fig. 1c,d**). Two independent clones of hPSCs carrying the deletion of TRPC6 (TRPC6 KO C21 and C47) were independently differentiated to cortical neurons. Loss of TRPC6 protein expression was confirmed by Western blot (**Fig. 1e**). TRPC6 KO and control hPSC-derived cortical neurons at 8 weeks post-differentiation did not show any significant differences in the expression levels of cortical neuron marker proteins (**Fig. 1b**).

### Functional characterization of hPSC-derived cortical neurons

Next, we validated the functional maturation of hPSC-derived cortical neurons using the whole-cell patch-clamp technique (**Fig. 2a-d**). Generation of action potential (AP) is the hallmark of neuronal differentiation and maturity of neurons given that presynaptic neurons generate and transmit AP to communicate with the postsynaptic neurons. AP was monitored in the current-clamp mode (**Fig. 2b**) and distributions of AP generation were analyzed. The frequency of “no AP”, “single AP”, or “multiple AP” was assessed in hPSC-derived cortical neurons at 0, 3, 6, or 8 weeks of differentiation (**Fig. 2c**). Cortical neurons differentiated for 6 and 8 weeks generated multiple and repetitive action potentials: ∼70% multiple AP and ∼30% single AP (**Fig. 2c**), whereas NPCs (0 week post-differentiation) had no multiple AP. Typical Na^+^ influx followed by K^+^ efflux upon membrane depolarization was observed in a voltage clamp mode from hPSC-derived cortical neurons after 6 weeks of differentiation (**Fig. 2d, Supplementary Fig. 3**).

**Figure 2.**
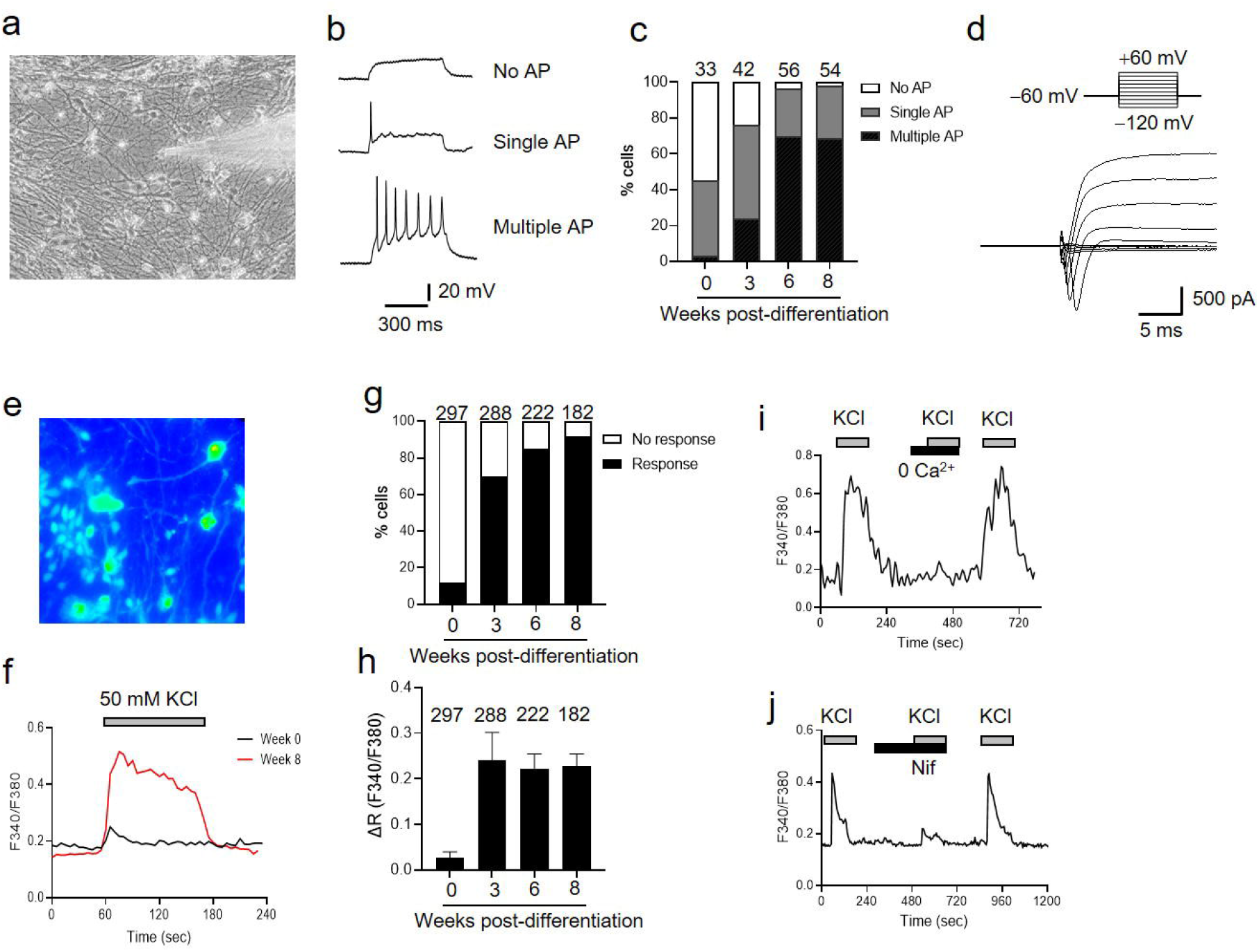
Electrophysiological characterization of hPSC-derived cortical neurons. (**a**) DIC image of hPSC-derived cortical neurons for electrophysiological whole-cell patch-clamp recording after 6 weeks of differentiation. (**b**) Whole-cell patch-clamp recording to monitor AP generated by injection of current pulses in a current clamp mode; no AP, single AP, or multiple and repetitive AP. (**c**) Distributions of AP generation; no AP, single AP, or multiple/repetitive AP in hPSC-derived cortical neurons differentiated for 0, 3, 6, and 8 weeks. Number of cells tested are shown from 3∼6 independent differentiation. (**d**) Na^+^ influx is followed by K^+^ efflux in a voltage clamp mode. (**e-j**) Characterization of calcium influx through VGCCs in hPSC-derived cortical neurons. (**e**) Image of Fura-2-loaded hPSC-derived cortical neurons after 6 weeks of differentiation. (**f**) Representative traces of intracellular calcium ions (Fura-2 F340/F380 ratio) in neurons stimulated by 50 mM KCl for 2 min; 0 week (black line) and 8 weeks (red line) of differentiation. (**g**) Percentage of neurons that evoke calcium influx by 50 mM KCl. Number of cells tested are shown from 3∼4 independent differentiation. (**h**) Net changes of calcium increase by 50 mM KCl. Data are means ± SEM from 3∼4 independent differentiation. (**i,j**) Representative calcium trace of Fura-2 F340/F380 ratio in 8-week old hPSC-derived cortical neurons in the absence of extracellular calcium ions (**i**) and in the presence of 2 μM nifedipine (Nif) (**j**).

We further examined functional activity of hPSC-derived cortical neurons using single-cell calcium imaging with a Fura-2 ratiometric calcium indicator (**Fig. 2e-j**). Neurons express voltage-gated calcium channels (VGCCs) that mediate calcium influx to trigger vesicle fusion and neurotransmitter release. Activity of VGCCs is a useful benchmark of neuronal differentiation and maturity of hPSC-derived cortical neurons. Upon applying 50 mM KCl to depolarize the membrane potential and specifically activate VGCCs, we analyzed calcium influx through VGCCs (**Fig. 2e,f**). Most cortical neurons (> 90%) after 6 and 8 weeks of differentiation evoked calcium influx upon 50 mM KCl stimulation (**Fig. 2g**) confirming that the efficiency of neuronal differentiation reaches 90∼100% (**Fig. 1b**). Net increase of calcium influx in a single neuron showed no differences between 3, 6, and 8 weeks of differentiation (**Fig. 2h**), however the percentage of neurons responding to KCl stimulation increased as cortical neurons matured (**Fig. 2g**). As a control, buffer without extracellular calcium ions (0 Ca^2+^) caused no calcium influx, and nifedipine, a L-type VGCC blocker, dramatically inhibited calcium influx (**Fig. 2i,j**), correlating with the prominent L-type VGCC in primary cortical neurons^24, 25^. Altogether, hPSC-derived cortical neurons after 6 and 8 weeks of differentiation were functionally active, whereas 3 week-old cortical neurons were relatively immature based on low percentages of neurons that generated multiple AP and responded to KCl stimulation (**Fig. 2c,g**).

### SOCE calcium signaling in mature hPSC-derived cortical neurons

SOCE modulates neuronal signaling and is associated with neurological diseases^7, 8, 9^ ^12, 17^. However, it remains controversial whether SOCE is active in primary neurons as recording SOCE in primary neurons is challenging due to intrinsic limitations^26^. We took advantage of hPSC-derived cortical neurons to monitor SOCE activity as neurons develop *in vitro*. Thapsigargin (TG), a SERCA pump inhibitor, depletes ER calcium stores and leads to activation of SOCE (**Fig. 3a**). TG-induced SOCE was weak in NPCs (0 week post-differentiation) but significantly strengthened as hPSC-derived cortical neurons matured over 8 weeks of differentiation (**Fig. 3b**). As a positive control of single-cell calcium imaging, ionomycin, a calcium ionophore, was applied to induce calcium influx in the presence of 2 mM CaCl_2_. Ionomycin caused consistent calcium influx, confirming the reliability and reproducibility of our calcium imaging experiments (**Fig. 3c,d**).

**Figure 3.**
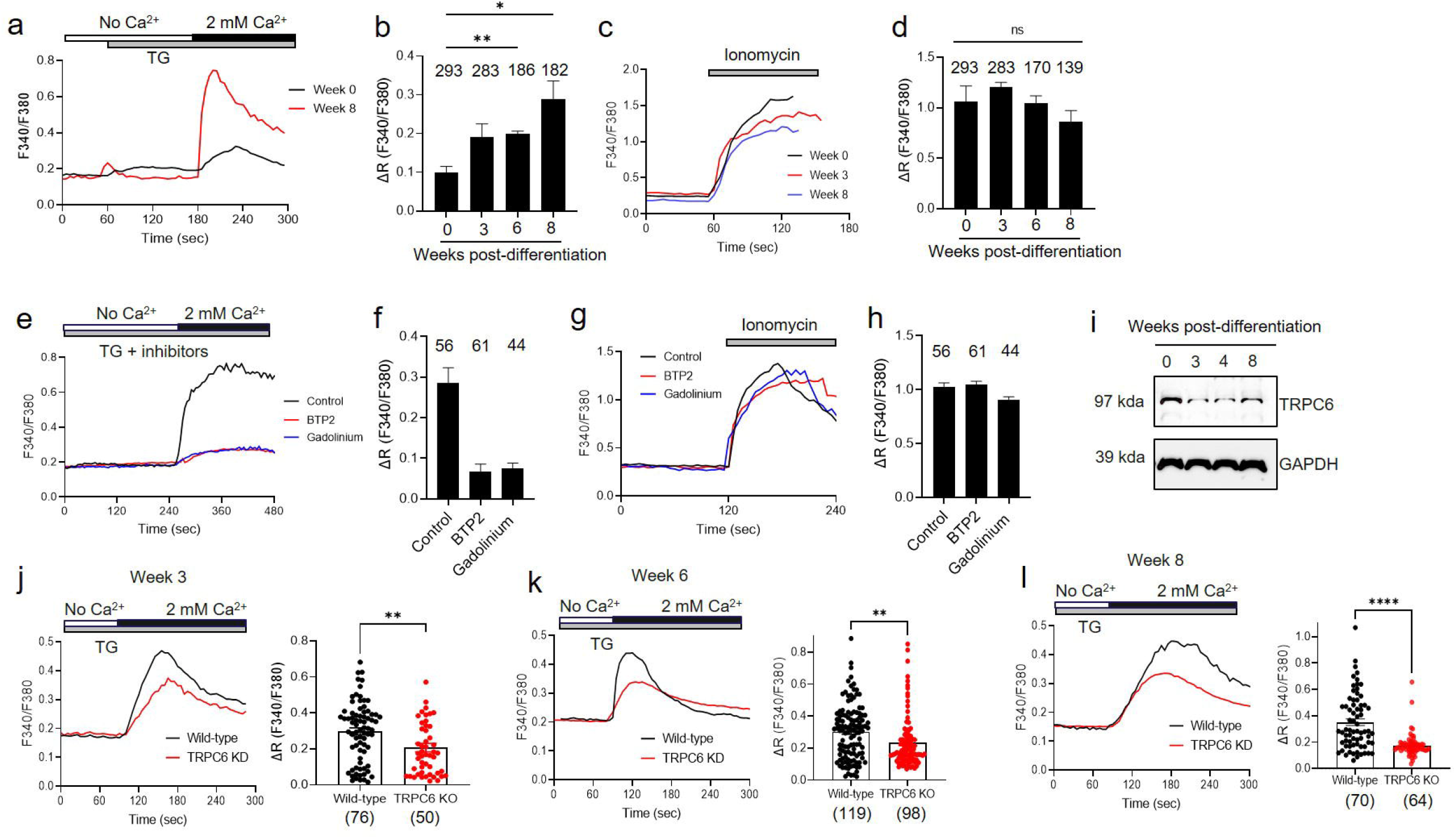
Downregulation of SOCE in TRPC6 KO hPSC-derived cortical neurons. (**a**) Representative calcium trace of Fura-2 F340/F380 ratio for SOCE responses by TG that depletes ER calcium store and leads to activation of SOCE; 0 week (black line) and 8 weeks (red line) of differentiation. (**b**) The net increase of calcium level by SOCE activation in hPSC-derived cortical neurons differentiated for 0, 3, 6, and 8 weeks. Data are means ± SEM and number of cells tested are shown from 5∼6 independent differentiation. (**c**) Representative calcium trace of Fura-2 F340/F380 ratio in ionomycin-treated cortical neurons. (**d**) The net increase of calcium level by ionomycin treatment as a control. Data are means ± SEM and number of cells tested are shown from 5∼6 independent differentiation. Welch and Brown-Forsythe one-way ANOVA was used in **b,d**. *, *p* < 0.05. **, *p* < 0.01. ns, not significant. (**e,f**) Representative calcium trace of SOCE in 8-week old cortical neurons in the presence of 10 μM BTP2, a selective TRPC inhibitor, or 5 μM gadolinium (Gd^3+^), a non-selective cation channel blocker. (**g,h**) Calcium increase in ionomycin-treated cortical neurons in the presence of BTP2 or Gd^3+^. Data in **f,h** are means ± SEM and number of cells tested are shown from 2 independent differentiation. (**i**) TRPC6 expression in hPSC-derived cortical neurons differentiated for 0, 3, 4, and 8 weeks. (**j-l**) TRPC6 KO reduces SOCE. Representative calcium trace of Fura-2 F340/F380 ratio (left) and quantification of net calcium increase (right) in hPSC-derived (C21 hPSC line) cortical neurons differentiated for 3 (**j**), 6 (**k**), and 8 (**l**) weeks. Data in **j-l** are means ± SEM and number of cells tested are shown in parentheses from 2 independent differentiation and unpaired two-tailed t-test was used. **, *p* < 0.01. ****, *p* < 0.0001.

Next, we tested SOCE using a pharmacological inhibitor, BTP2, a selective TRPC channel blocker without subtype selectivity ^27, 28^. BTP2 treatment strongly inhibited SOCE in 8-week old hPSC-derived cortical neurons, comparable to gadolinium (Gd^3+^), a non-selective cation channel blocker^29^ (**Fig. 3e,f**). Both BTP2 and gadolinium had no effect on ionomycin-induced calcium influx (**Fig. 3g,h**), indicating that TRPC channels are linked to SOCE.

TRPC6 is involved in SOCE^18^ and human TRPC6 is expressed throughout the CNS and peripheral tissues^30^. We observed TRPC6 expression in hPSC-derived cortical neurons during neuronal differentiation (**Fig. 3i**). To validate that TRPC6 regulates SOCE, we tested SOCE in TRPC6 KO hPSC-derived cortical neurons (C21 hPSC line) (**Fig. 3j-l**). Indeed, TRPC6 KO reduced SOCE in cortical neurons at 3 (**Fig. 3j**), 6 (**Fig. 3k**), and 8 weeks (**Fig. 3l**) of differentiation. We confirmed the inhibition of SOCE by TRPC6 KO using different hPSC line (C47)(**Supplementary Fig. 4a**). Altogether, our data support that SOCE is impaired in TRPC6 KO neurons, which recapitulate ASD pathology.

### Hyperexcitability in TRPC6 KO hPSC-derived cortical neurons

Next, we performed whole-cell patch-clamping to analyze neuronal activity in TRPC6 KO cortical neurons, where SOCE is reduced. Intriguingly, TRPC6 KO increased the frequency of AP generation in hPSC-derived cortical neurons (C21 hPSC line) differentiated for 6 weeks (**Fig. 4a-d**) and 8 weeks (**Fig. 4e,f**). All TRPC6 KO neurons produced multiple AP, whereas approximately 30% and 20% of wild-type cortical neurons generated single AP in week 6-old and week 8-old neurons, respectively (**Fig. 4c,e**). This increase of AP frequency was confirmed in different hPSC line (C47)-derived TRPC6 KO cortical neurons (**Supplementary Fig. 4b-d**).

**Figure 4.**
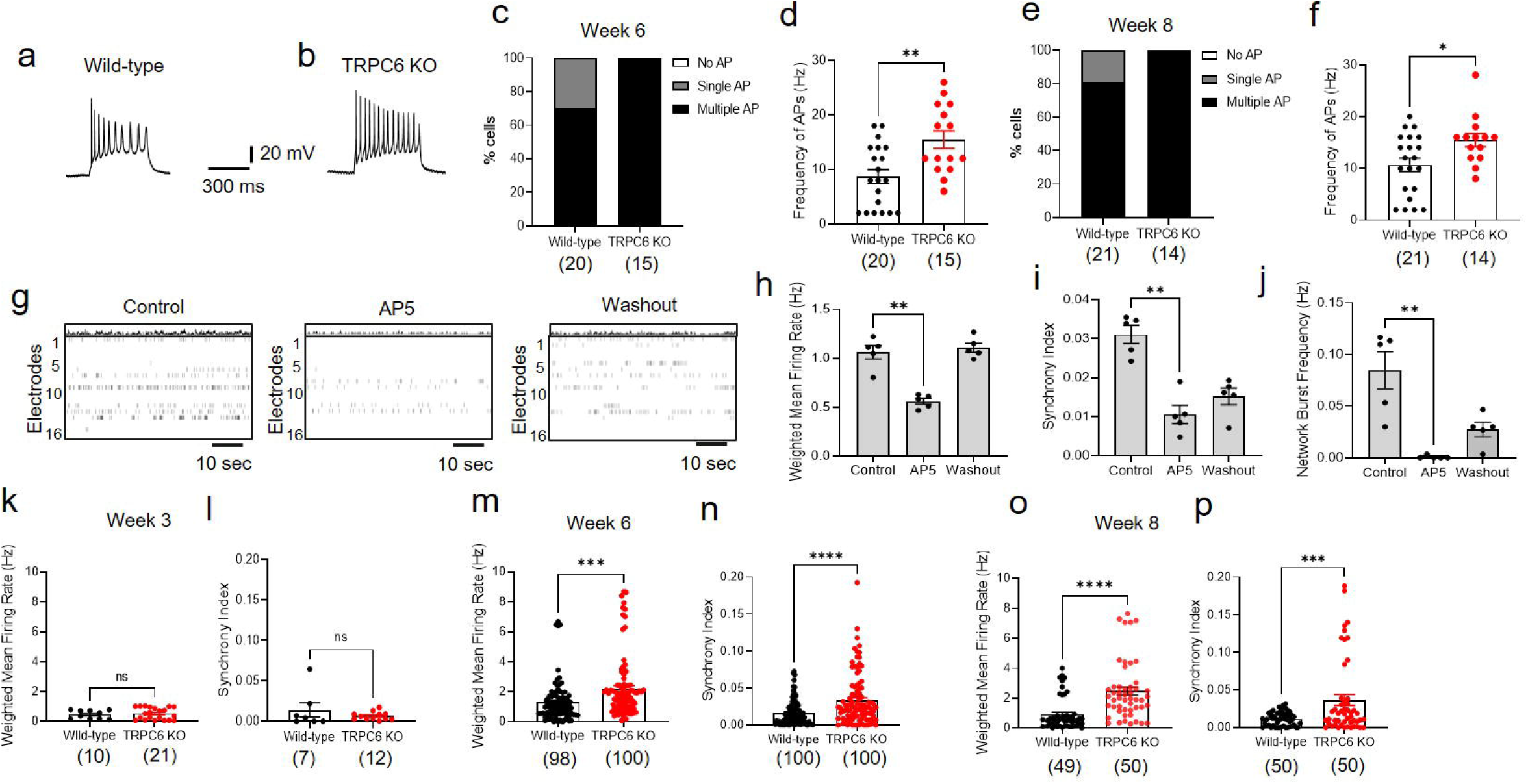
Hyperexcitability in TRPC6 KO hPSC-derived cortical neurons. (**a,b**) Representative multiple AP of wild-type and TRPC6 KO neurons differentiated for 6 weeks (C21 hPSC line). APs were generated by injection of current pulses in a current clamp mode. (**c**) Distributions of AP generation of either no AP, single AP, or multiple/repetitive AP and (**d**) frequency of APs in wild-type and TRPC6 KO hPSC-derived cortical neurons differentiated for 6 weeks and 8 weeks (**e,f**). (**g-p**) Monitoring neuronal activity and neural network using microelectrode array (MEA). (**g**) Raster plots of hPSC-derived cortical neurons at 8 weeks of differentiation in the presence of 1 μM AP5 (5 min) and recovery after washout. Quantification of weighted mean firing rate (Hz) (**h**), synchrony index (**i**), and network burst frequency of neural network (**j**). Data are means ± SEM (n = 5) from 2 independent differentiation. (**k-p**) TRPC6 KO increases neuronal activity and neural network formation (C21 hPSC line). Quantification of weighted mean firing rate (Hz), synchrony index of neural network from wild-type and TRPC6 KO hPSC-derived cortical neurons differentiated for 3 (**k,l**), 6 (**m,n**), and 8 (**o,p**) weeks. Data in **d,f,k-p** are means ± SEM and number of cells tested are shown in parentheses from 2 independent differentiation and unpaired two-tailed t-test was used. *, *p* < 0.05. **, *p* < 0.01. ***, *p* < 0.001. ****, *p* < 0.0001.

We further tested neuronal activity and neural network in TRPC6 KO hPSC-derived cortical neurons using micro-electrode array (MEA). MEA contains a grid of 16 tightly spaced electrodes embedded in the culture surface seeded with hPSC-derived cortical neurons (**Fig. 4g**). MEA simultaneously monitors neuronal activity from different locations across the cultured cortical neurons to detect propagation and synchronization of neural activity, thus measuring both the neuronal activity and neural network formation. As a control, we confirmed that tetrodotoxin (TTX), a selective voltage-gated sodium channel blocker inhibiting AP generation, completely blocked weighted mean firing rate and synchrony, which were rescued by washout (**Supplementary Fig. 5**). In addition, amino-phosphonopentanoate (AP5), a selective NMDA receptor antagonist, inhibited not only weighted mean firing rate, but also synchrony and network bursts frequency (**Fig. 4g-j**), indicating that synaptic transmission and neural networking in hPSC-derived cortical neurons are mainly mediated by glutamatergic excitatory synapses. Correlating with AP generation (**Fig. 4a-f**), TRPC6 KO led to an increase of weighted mean firing rate and synchrony in hPSC-derived cortical neurons at 6 and 8 weeks of differentiation (**Fig. 4m-p**), however TRPC6 KO had little effect on neuronal activity of 3-week old immature cortical neurons (**Fig. 4k,l**). Taken together, electrophysiological data show that TRPC6 KO hPSC-derived cortical neurons have hyperexcitability of neuronal activity and neural network.

### Transcriptome profiling of TRPC6 KO hPSC-derived cortical neurons

To gain insight to the molecular basis of the changes in electrophysiological activity, we applied whole transcriptome RNA-Seq analysis to characterize gene expression differences in cortical neurons derived from control and TRPC6 KO hPSC at 8 weeks of differentiation. RNA-Seq was performed on four independent differentiation experiments; biological replicates of control and TRPC6 KO hPSC-derived cortical neurons differentiated from two different TRPC6 KO hPSC clones (C21 and C47). We categorized upregulated and downregulated genes with at least 1.5 fold change (FC) and < 0.05 *p*-value cut-off: 362 upregulated and 132 downregulated transcripts were identified (**Fig. 5a, Supplementary Table 1**). The hierarchical clustering based on differentially expressed RNA transcripts revealed clear clustering of four biological replicates in each condition (**Fig. 5b**). Gene ontology (GO) enrichment analysis revealed that the most significantly upregulated biological processes in TRPC6 KO hPSC-derived cortical neurons include ‘chemical synaptic transmission’, ‘trans-synaptic signaling’, and ‘synapse organization’ (**Fig. 5c**); note that no biological processes significantly downregulated were observed. We further applied KEGG pathway analysis, showing the upregulation of ‘glutamate signaling’, ‘calcium signaling’, and ‘neurotransmitter receptor activity’ (**Supplementary Fig. 6**). vGLUT1 mRNA was upregulated, whereas vesicular GABA transporter (VGAT) mRNA was downregulated (**Supplementary Fig. 6a,b**). Furthermore, mRNA levels of calcium signaling pathways, presynaptic proteins, glutamate ionotropic receptor kainate type subunit 1 (GRIK1), and glutamate metabotropic receptor (GRM1) were significantly elevated in TRPC6 KO hPSC-derived cortical neurons (**Supplementary Fig. 6a,c,d and Supplementary Table 1**), thereby leading to the hyperactivity of TRPC6 KO neurons. Altogether, our transcriptomic analyses support the hyperactivity of TRPC6 KO hPSC-derived cortical neurons by upregulating glutamate excitatory synapses and calcium signaling pathways.

**Figure 5.**
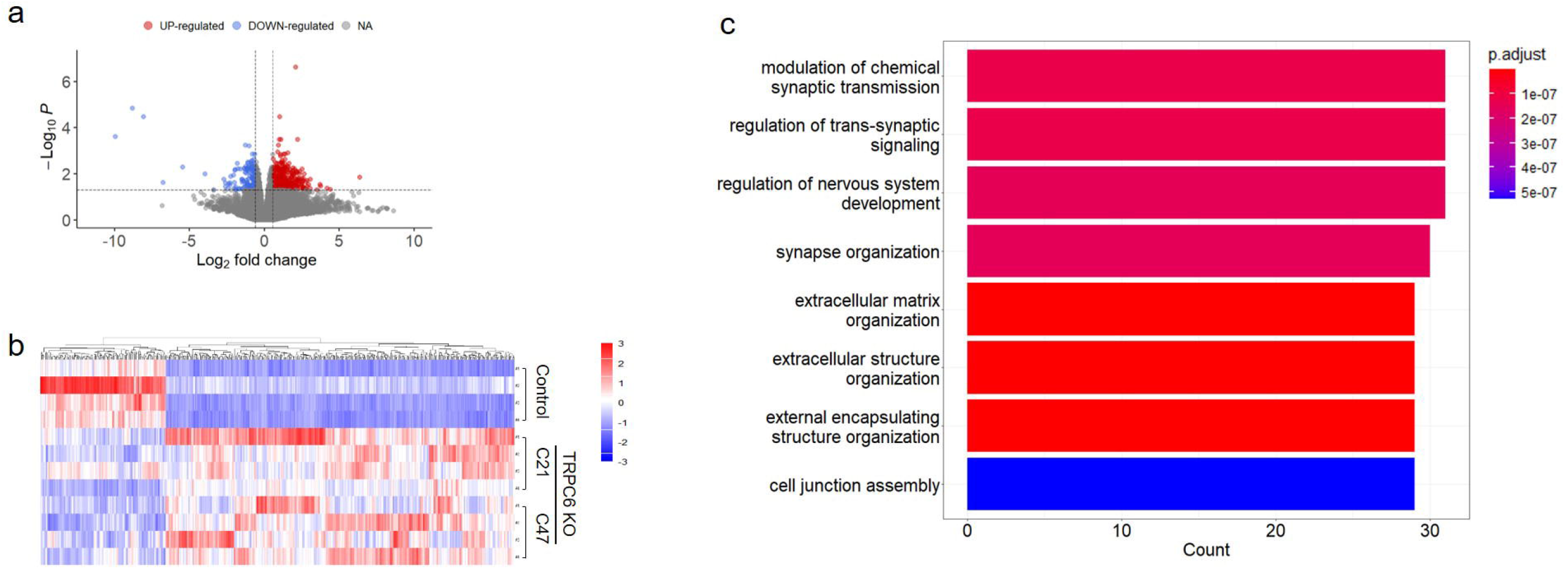
Transcriptome profiling of TRPC6 KO hPSC-derived cortical neurons. (**a**) Volcano plot of the differentially expressed (DE) mRNAs between control and TRPC6 KO hPSC-derived cortical neurons at 8 weeks of differentiation; upregulated (light red) and downregulated (light blue) transcripts. Total 27,814 mRNA transcripts were tested. Color indicates significantly dysregulated transcripts with < 0.05 *p*-value and 1.5 fold change (FC) cut-off. (**b**) Heatmap of hierarchical clustering analysis of DE mRNAs representing the upregulation and downregulation of transcripts from three different conditions; control hPSC-derived cortical neurons and TRPC6 KO hPSC-derived cortical neurons from two different hPSC clones (C21 and C47) at 8 weeks of differentiation. Four biological replicates and independent differentiation were analyzed (1.5 FC, *p* < 0.05). Red color, upregulated mRNAs; blue color, downregulated mRNAs. Expression data have been standardized as *z*-scores for each mRNA. (**c**) GO enrichment analysis for biological process of upregulated genes in TRPC6 KO hPSC-derived cortical neurons. The GO cut-off criteria included *q* (adjusted *p* value) < 0.000001.

## Discussion

Neurological and psychiatric disorders are often caused by dysregulation of synaptic transmission and neural networks. Given that the access to the human brain is almost impossible, studying human neural networks in disease conditions is challenging due to the absence of good model system. Most studies have used various animal models that have limitation to reproduce human brain development and human neurophysiology. Patient-specific hPSC-derived neurons with human genetic and epigenetic backgrounds recapitulate the genomic, molecular and cellular properties of native human neurons thereby offering a human cell-based model to investigate the pathophysiology. We have exploited this model system to demonstrate that TRPC6 KO hPSC-derived cortical neurons reproduce hyperexcitability of ASD phenotype and study key aspects of ASD pathology.

ASD is a highly heterogeneous neurodevelopmental disorder. ADHD and hyperactivity are common comorbidities of ASD: >50% ASD individuals have ADHD^31, 32^. Due to the complexity, it remains challenging to unveil the pathophysiological and molecular mechanisms underlying this hyperactivity and hyperexcitability of neurons in ASD. Using TRPC6 KO hPSC-derived cortical neurons we can reproduce hyperexcitability of ASD phenotype. TRPC6 KO reduces SOCE in cortical neurons after 3 to 8 weeks of differentiation (**Fig. 3j-l**). Hyperexcitability is only observable in mature TRPC6 KO cortical neurons differentiated for 6 to 8 weeks (**Fig. 4a-p**), not in 3-week old immature cortical neurons (**Fig. 4k,l**). RNA-Seq analysis further validates upregulation of excitatory synapses and downregulation of inhibitory GABA transporter (VGAT), implying imbalance of synaptic connections and networks that may result in hyperexcitability of TRPC6 KO hPSC-derived cortical neurons (**Fig. 5**).

Our data provide evidence that reduction of SOCE by TRPC6 KO might result in hyperexcitability of mature cortical neurons (**Fig. 4m-p**). TRPC6 loss-of-function mutations in *Drosophila* cause ASD-like behavior including hyperactivity^21^, suggesting that our TRPC6 KO hPSC-derived cortical neurons can be a good model to understand hyperactive behavior of ASD at a cellular and molecular level. These hyperactive hPSC-derived cortical neurons will be used for better care of ASD by taking a personalized medicine approach. Every ASD individuals show different and heterogeneous pathophysiology. We can apply different therapeutics depending on different pathology of hPSC-derived cortical neurons differentiated from every ASD individual, taking into account individual variability in genetics and pathophysiology.

Despite the significance of SOCE in cellular signaling, it has been challenging to monitor neuronal SOCE in primary neurons^26, 33^. SOCE is very weak in hippocampal neurons^33^, so neuronal SOCE has remained controversial to exist^26^. We took advantages of hPSC-derived cortical neurons to monitor neuronal SOCE during development over one time point. We show that neuronal SOCE activity becomes stronger and significant in mature hPSC-derived cortical neurons as a neuronal SOCE model system, and TRPC6 is involved in SOCE pathway.

TRPC6 has high selectivity for Ca^2+^ relative to Na^+^ and contributes to SOCE upon ER calcium store depletion^34, 35, 36, 37^. Disruption of the TRPC6 gene may contribute to the ASD phenotype^19^ and TRPC6 loss-of-function mutations result in hyperactivity in *Drosophila*^21^. TRPC channels including TRPC6 can be therapeutic targets for intervention of ASD. Therefore, our TRPC6 KO hPSC-derived cortical neurons provide a platform to screen therapeutics that rescue SOCE and reverse hyperexcitability. It remains a topic of further study if TRPC6 and TRPC agonists can rescue SOCE and reverse hyperactive phenotype as ASD therapeutics.

## Methods

### Reagents

Fura-2 pentaacetoxymethyl ester (fura-2/AM) was from Thermo Fisher Scientific (Waltham, MA, USA).

### CRISPR/Cas9 editing and PSCs maintenance

The hPSC lines used in this study is CRTD5^38^ generated from BJ fibroblasts (CRL-2522, ATCC). Cells were cultured and maintained on Matrigel-coated (BD bioscience, cat# 354277) plates in mTeSR1 medium (STEM CELL technologies, cat# #85850) in a humidified incubator at 37°C and 5% CO_2_. For editing, the guide RNA (gRNA) sequence targeting the first exon of TRPC6 was selected using CRISPR-Cas9 guide RNA design tool (Intergrated DNA technologies). Single guide RNA (sgRNA) was synthesized using EnGen sgRNA Synthesis Kit (NEB, E3322) according to the manufacturer’s instructions. Nucleofection was carried out using the Amaxa nulceofection system (P3 primarycell 4D-nucleofector kit, Cat#V4XP-3032) according to the manufacturer’s instructions. Briefly, RNP complex were generated by mixing 1 μg of sgRNA with 2 μM of EnGen SpyCas9 NLS (NEB, M0646) at room temperature for 15-20 min. 2.5-3 × 10^5^ hPSCs were electroporated using CB150 nucleofection program and plated onto Matrigel-coated plates. After 48 hr, the cells were diluted and plated as a single cell on Matrigel-coated plates for 10-15 days to make colonies. Genomic DNA (gDNA) was extracted using quick extract genomic DNA extraction buffer (epicenter). The region of TRPC6 targeted by sgRNA was amplified with specific primers (Forward: TGTTGACATAGTAACTCTTCAGCTCCGTCTCCCTTGC, Reverse: GCTGCCTTGCTACGGCTACTACCCCT) and sequenced using PCR-Master mix (Thermo Fisher Scientific).

### Stem cell differentiation into cortical neurons

hPSCs were differentiated into cortical neurons following previously published protocol^23^, with minor modifications. To initiate differentiation, hPSC colonies were dissociated into single cells using TrypLE (Thermo Fischer Scientific) and plated onto Matrigel-coated (BD Biosciences) plates in mTeSR1 medium (Stem cell Technologies, Vancouver) containing 10 μM Y-276321 (ROCK inhibitor). Next day, the cells were 90-100% confluent and differentiation was initiated by changing medium to Neurobasal medium (DMEM/F12, Neurobasal, 1X B-27 minus vitamin A, 1X N2 supplement, 1X L-Glutamine, 1X Non-essential amino acids (NEAA), 50 μM β-mercapto-ethanol, 0.2X Penicillin/streptomycin) supplemented with 10 μM SB431542 and 2 μM Dorsomorphin for 12 days. At day 10, the cells were split using TrypLE and plated onto Matrigel-coated plates in neurobasal media containing 5 μM Rock inhibitor. For neural proliferation (days 14-18), the neurobasal media was supplemented with 20 ng/ml bFGF. At day 20, NPCs were cryopreserved or plated for maturation onto Matrigel-coated plates in neurobasal media supplemented with 10 ng/ml BDNF, 10 ng/ml GDNF, 2 μg/ml insulin, 20 μM dibutyryl-cyclic AMP (db-cAMP, Sigma), and 200 μM Ascorbic acid (AA, Sigma). At day 28, the cells were plated for experiment at a density of 50,000 cells/cm^2^ onto 100 μg/ml poly-L-ornithine (PO, Sigma) and 20μg/ml laminin-coated plates and media was changed every 2-3 days. The cells were matured for 6-8 weeks.

### Immunostaining

Cells were washed with 1X PBS and fixed with 4% paraformaldehyde for 15 min at room temperature. The fixed cells were washed three times with PBS, treated with 0.2 % Triton X-100 (Sigma-Aldrich) in PBS for 30 min and blocked in PBST (PBS with 0.2% tween-20) containing 3% bovine serum albumin (BSA) for 2-3 hr. The cells were incubated with primary antibodies overnight at 4°C. Primary antibodies consisted of SOX2 (Rabbit, 1:200, Invitrogen: MA1-014), FOXG1 (Rabbit, 1:200, Abcam: ab18259), OTX2 (Goat, 1:300, R&D: AF1979), Nestin (Mouse, 1:100, Invitrogen: MA1110, MAP2 (chicken, 1:500, Abcam: ab5392), MAP2 (Mouse, 1:500, Invitrogen: 13-1500), beta tubulin III (Mouse, 1:300, MAB1637), TBR2 (Rabbit,1:300, Cell signaling: 66325), CTIP2 (Rabbit, 1:200, Cell signaling:), BRN2 (Rabbit, 1:200, Cell signaling:12137), GFAP (Chicken, 1:400, Abcam: ab4674), and SATB2 (Rabbit, 1:200, Invitrogen:PA5-83092). Next day, the cells were washed three times with PBST at 10 min intervals and incubated with the secondary antibodies diluted 1:1000 in PBST containing 3% BSA for 1 hr at room temperature. Secondary antibodies were conjugated with Alexa Flour 488, Alexa Flour 555, and Alexa Flour 647 dyes (all Thermo Fischer Scientific). Nuclei were stained with DAPI (Thermo Fischer Scientific) for 5 min. Cells were washed three times with PBS and imaged using the inverted fluorescence microscope (Olympus IX 53).

### RNA extraction, Quantitative PCR, and library preparation

The cells were lysed in TRIzol (Thermo Fischer Scientific) and the total RNA was extracted with Direct-zol RNA extraction kit (Zymo Research) following the manufacturer’s instructions. Complementary DNA was synthesized from 500 ng of RNA using RevertAid First Strand cDNA Synthesis kit (Thermo Fischer Scientific). Quantitative PCR (qPCR) was performed using Syber Green PCR Master Mix (Applied biosystems) with the primers listed in **Supplementary Table 2**. For library preparation, total RNA with a RNA integrity number (RIN) above 8 was used as input using TruSeq Stranded mRNA kit (Cat #: 20020594) from Illumina following the manufacturer’s protocol. Briefly, from 500 ng of total RNA, mRNA molecules were purified using poly-T oligo attached magnetic beads and then mRNA was fragmented. cDNA was generated from the cleaved RNA fragments using random priming during first and second strand synthesis. Barcoded DNA adapters was ligated to both ends of DNA, and then amplified. The quality of library generated was checked on an Agilent 2100 Bioanalyzer system and quantified using a Qubit system. Libraries that pass quality control was pooled, clustered on a cBot platform, and sequenced on an Illumina HiSeq 4000 at a minimum of 20 million paired end reads (2×75 bp) per sample.

### RNA-Seq data analysis

Starting with the FASTQ files, trimming, aligning, and transcript quantification were performed within the Galaxy platform^39^. The paired-end reads were trimmed with default parameter settings using Cutadapt. Alignment of the reads to reference genome GRCh38/hg38 was carried out using HiSAT2^40^. Transcript counting was performed with featureCounts^41^. The count matrix was then normalized and differential expression analysis was performed with the R-library EdgeR^42^. *P*-values were adjusted with the Benjamini–Hochberg procedure, which controlled the false discovery rate (FDR). Genes with adjusted *p* values < 0.05 and fold changes > 1.5 were considered to be differentially expressed. The volcano plot and heatmap were created using the EnhancedVolcano and Pheatmap R-libraries, respectively. The heatmap was z-score scaled column wise; a *z*-score normalization was performed on the normalized read counts across samples for each gene. *Z*-scores were computed to plot a heatmap on a gene-by-gene basis by subtracting the mean and then dividing by the standard deviation. Gene Ontology (GO) enrichment and KEGG pathway analysis were performed using the Over-Representation Analysis (ORA) functions of the clusterProfiler 4.0 R-library^43^. Pathview graphs of differentially expressed genes on KEGG pathways was performed using the Pathview R-library^44^.

### Electrophysiology

The action potentials were recorded using the whole-cell patch-clamp technique using an EPC-10 USB amplifier (HEKA Elektronik, Lambrecht/Pfalz, Germany). Data acquisition, voltage control, and analysis were accomplished using software (HEKA Patchmaster). Neurons were placed in the chamber and perfused with the normal Tyrode’s bath solution (mM): 143 NaCl, 5.4 KCl, 0.33 NaH2PO4, 0.5 MgCl2, 5 HEPES, 2 CaCl2, and 11 glucose; pH 7.4 adjusted with NaOH. Patch pipettes were pulled from borosilicate capillary tubes (A-M systems, WA, USA) using a puller PC-10 (Narishige, Tokyo, Japan) and filled with an internal solution (mM): 130 K-gluconate, 3 KCl, 2 MgCl2, 10 HEPES, 5 Na2ATP, 0.5 Na2GTP, 0.2 EGTA; pH 7.3 adjusted with KOH. The resistance of patch pipettes was 3∼5 MΩ. Action potentials were generated by a series of current steps from −20 to +60 pA for 500 ms in a current-clamp mode. Whole-cell currents were measured by a series of 20 mV voltage steps from −120 to +60 mV for 1 sec in a voltage-clamp mode. Signals were low-pass filtered with a cut-off frequency of 5 kHz and sampled at 10 kHz.

### Multi-well microelectrode array (MEA) analysis

48-well MEA plates were coated with poly-L-ornithine and laminin for neuronal differentiation as described before. NPCs (15,000 cells/well) were plated in the center of the well. Extracellular recordings were carried out using Axion’s Maestro multi-well 768 electrode recording platform in combination with Axion 48-well MEA plates (Axion Biosystems). Each well contains a 4×4 16 channel electrode array with four reference electrodes. Extracellular voltage recordings were collected at a sampling rate of 12.5 kHz per channel. A band-pass filter (200 Hz to 3 kHz cut-off frequencies) was applied with a variable threshold spike detector at ±6 standard deviations of the root mean square (RMS) of the background noise.

Recordings were performed before media change or two days after media change. During recording, the MEA plate was maintained at 37°C and 5% CO2. MEA data analysis was performed using the Axion Biosystems Neural Metric Tool (Axion Biosystems). An electrode was considered active at a threshold of 3 spikes/min. Network Bursts was defined as at least 5 consecutive spikes across multiple electrode with interspike intervals (ISI) of less than 100 ms and a minimum of 25% electrodes.

### Calcium imaging

The neurons on the coverslip were loaded with 3 μM Fura-2AM (Thermo Fisher Scientific) for 30 min at room temperature. Calcium imaging experiments were carried out using a monochromator-based spectrofluorometric system (Photon Technology International, Lawrenceville, NJ) with Evolve 512 camera (Teledyne Photometrics, AZ, USA). Dual excitation and emission were at 340/380 and 510 nm, respectively. Data acquisition was accomplished using EasyRatioPro software. Neurons were perfused with the normal Tyrode’s bath solution (mM): 143 NaCl, 5.4 KCl, 0.33 NaH2PO4, 0.5 MgCl2, 5 HEPES, 2 CaCl2, and 11 glucose; pH 7.4 adjusted with NaOH. 50 mM KCl was applied to depolarize the membrane potential to evoke calcium influx through voltage-gated calcium channels. Regions of interest (ROI) were assigned by highlighting the perimeter of the cell using the software. Mean fluorescence intensity was recorded within the ROIs. Changes in fluorescence intensity were analyzed after background subtraction using ImageJ software (National Institutes of Health, Bethesda, MD).

### Western Blot

Total proteins were extracted with the Laemmle sample buffer (200mM Tris-HCl, pH 6.8, 8% sodium dodecyl sulfate, 0.4% Bromophenol blue, 20% glycerol, 5% 2-mercaptoethanl) and heated at 95°C for 10 min. Cell lysates and protein samples were loaded on Bolt™ 4-12% Bis-Tris Plus Precast Protein Gels (invitrogen NW04122BOX), and proteins were transferred to nitrocellulose membrane (88018, Thermo Fisher Scientific). Blots were then blocked with 5% skim milk in TBST for at least 1 hr at room temperature. Immunoblotting was done overnight at 4°C with the following antibodies at the appropriate dilutions: TRPC6 antibody (Abcam ab228771, 1:1000). The blots were washed the next day and incubated with Goat anti-Rabbit-HRP secondary antibody (Cat # 31460, Thermo Fisher Scientific, 1:10,000). The Protein bands were subsequently scanned using the ChemiDoc imaging system (BioRad).

### Statistical analysis

Data analysis was performed using OriginPro 2019 software (OriginLab Corporation, Northampton, MA, USA) and GraphPad Prism 9 (GraphPad Software, San Diego, CA, USA). Data are means ± standard error of the mean (S.E.M.). Welch and Brown-Forsythe one-way ANOVA was used to determine any statistically significant differences between three or more independent groups. Unpaired two-tailed t-test was used to estimate statistical significance between two groups. Probabilities of *p* < 0.05 was considered significant.

## Acknowledgements

We thank Dr. Volker Busskamp for generous gift of the CRDT5 cell line. We thank Dr. Shahryar Khattak, Dr. Khaled Machaca and Dr. Raphael Courjaret for technical support. Thanks to Dr. Khalid Ouararhni for RNA-Seq. This work was supported by the grant from Qatar Biomedical Research Institute (Project Number SF 2019 004 to Y.P.).

## Supplementary Figure legend

**Supplementary Figure 1. TRPC6 KO in CRTD5 hPSC lines using CRISPR/Cas9**. (**a**) Genomic DNA sequence of TRPC6 intronic and exonic sequence spanning the start codon (green) and guide RNA targeting sequence. (**b,c**) Genomic DNA sequencing analysis of KO clones. PCR primers were designed to amplify the gRNA-targeted region. Sequence of KO clone 21 (65 bp deletion), clone 47 (49 bp deletion), and wild-type TRPC6. KO clones were identified by Sanger sequencing. (**d**) PCR products from the intact DNA sequence, 534 bp.

**Supplementary Figure 2**. (**a**) Bright field images of control (Ctrl) and KO clones (C21 and C47). (**b**) Expression of pluripotency markers; OCT4, NANOG, and SOX2. TRPC6 mRNA in KO clones. (**c**) Immunostaining of control NPCs and TRPC6 KO NPCs derived from two different hPSC clones (C21 and C47) with antibodies against SOX2, OTX2, FOXG1, and Nestin (Nes). Nuclei are stained with DAPI. Scale, 200 μm.

**Supplementary Figure 3**. Whole-cell currents in hPSC-derived cortical neurons differentiated for 6 weeks from NPCs were measured using whole-cell patch-clamp in a voltage-clamp mode. Ionic current is generated by a series of 20 mV voltage steps from −120 to +60 mV for 1 sec in a voltage-clamp mode. Na^+^ channels are rapidly inactivated, whereas K^+^ channels remain open upon membrane depolarization.

**Supplementary Figure 4**. (**a**) Reduced SOCE in different hPSC line (C47)-derived TRPC6 KO cortical neurons. Quantification of net calcium increase in hPSC-derived cortical neurons (C47 hPSC line) differentiated for 8 weeks. (**b**) Representative multiple AP of wild-type and different hPSC line (C47)-derived TRPC6 KO neurons differentiated for 6 weeks. (**c**) Distributions of AP generation of either no AP, single AP, or multiple/repetitive AP and (**d**) frequency of APs in wild-type and TRPC6 KO hPSC-derived cortical neurons (C47 hPSC line). Data in **a-c** are means ± SEM and number of cells tested are shown in parentheses from 2 independent differentiation and unpaired two-tailed t-test was used. *, *p* < 0.05. ***, *p* < 0.001.

**Supplementary Figure 5**. (**a**) Raster plots showing electrical activity of hPSC-derived cortical neurons at 8 weeks of differentiation in the presence of 1 μM TTX (5 min) and recovery after washout. Each row of spikes represents an electrode; 16 electrodes in a single well. Vertical red rectangles represent events of network bursts of electrical activity. Quantification of weighted mean firing rate (Hz) (**b**) and synchrony index (**c**). Data in **b,c** are means ± SEM (n = 5) from 2 independent differentiation.

**Supplementary Figure 6**. KEGG Pathway enrichment analysis using clusterProfiler and Pathview. KEGG view on (**a**) glutamatergic presynaptic neurons, (**b**) GABAergic presynaptic neurons, (**c**) calcium signaling pathway, and (**d**) synaptic vesicle cycle pathway. Colors in **a-d** correspond to log_2_ FC (fold changes) between control and TRPC6 KO hPSC-derived cortical neurons. Blue, downregulated; red, upregulated.

