## supplemental file for "Deletion of *TRPC6*, an autism risk gene, induces hyperexcitability in cortical neurons derived from human pluripotent stem cells"

### Supplementary Figure 1

a

5’CTGCCCACTCGGCTCCCCCGGGAGCGGGGCCCAGGCCAGTCGGGCGTTCCCGCCATGAGCCAGAGCCCGGCGTTCGGCCCCGGAGGGGCAGTTCTCCCCGG 3’
3’GACGGGTGAGCCGAGGGGGCCCTCGCCCCGGGTCCGGTCAGCCCGCAAGGGCGGTACTCGGTCTCGGGCCGCAAGCCCGGGGCCTCCCCGTCAAGAGGGGCC 5’
CAGCCCGCAAGGGCGGTACT (guide RNA)

b

5’CTGCCCACTCGGCTCCCCCGGGAGCGGGGCCCAGGCCAGTCGGGCGTTCCCGCCATGAGCCAGAGCCCGGCGTTCGGCCCCGGAGGGGCAGTTCTCCCCGG 3’ (Wild-type)
5’CTGCCCACTCGGCTCCCCCGGGAGC-----------------------------------------------------------------AGTTCTCCCCGG 3’ (TRPC6 KO, C21)
5’CTGCCCACTCGGCTCCCCCGGGAGCGGGGCCCAG-------------------------------------------------GAGGGGCAGTTCTCCCCGG 3’ (TRPC6 KO, C47)

c

TRPC6 KO (C21), 65 nucleotides deletion

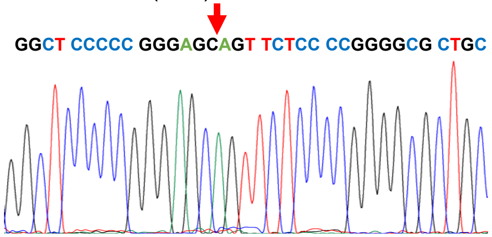


TRPC6 KO (C47), 49 nucleotides deletion

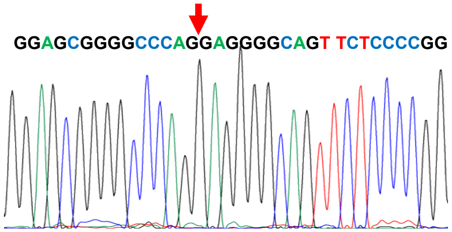


d

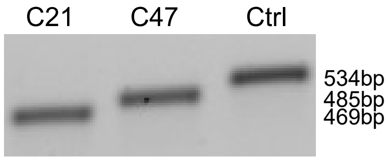

### Supplementary Figure 2

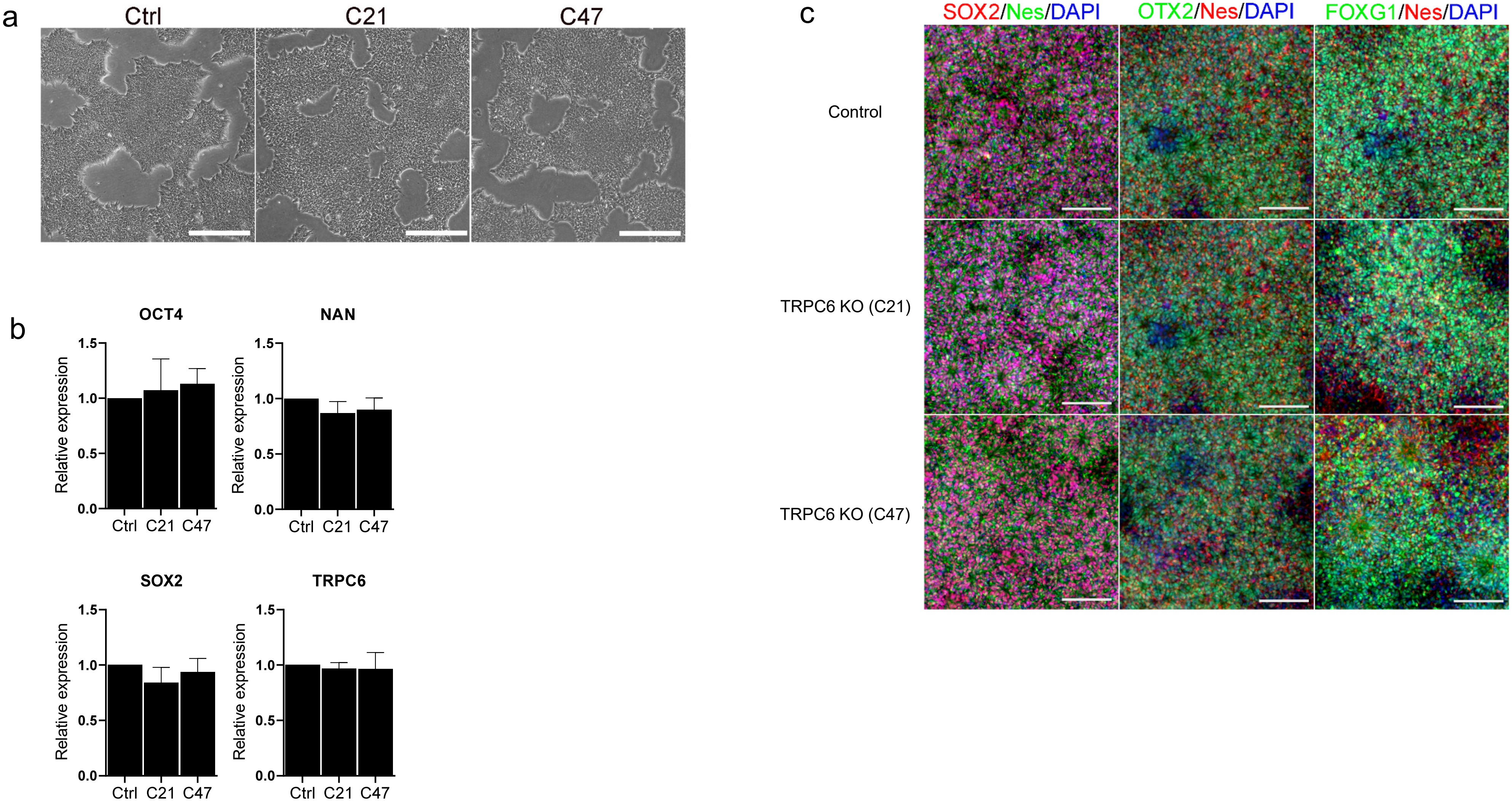

### Supplementary Figure 3

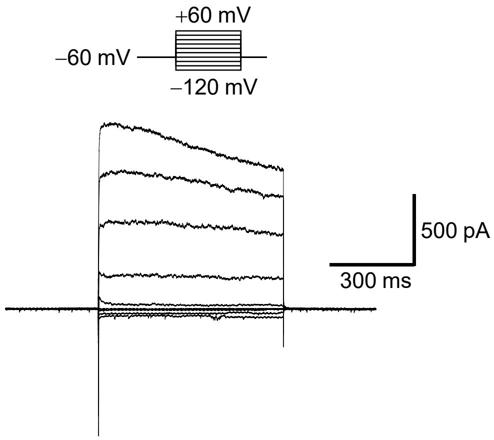

### Supplementary Figure 4

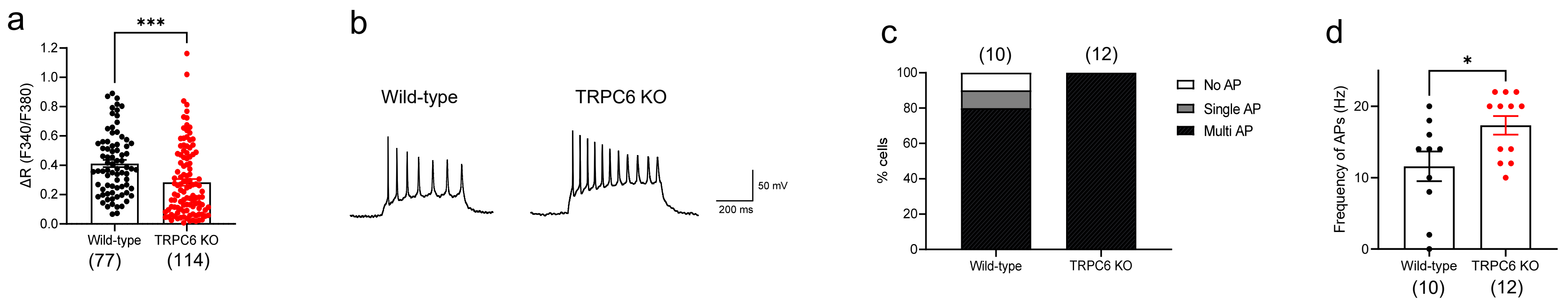

### Supplementary Figure 5

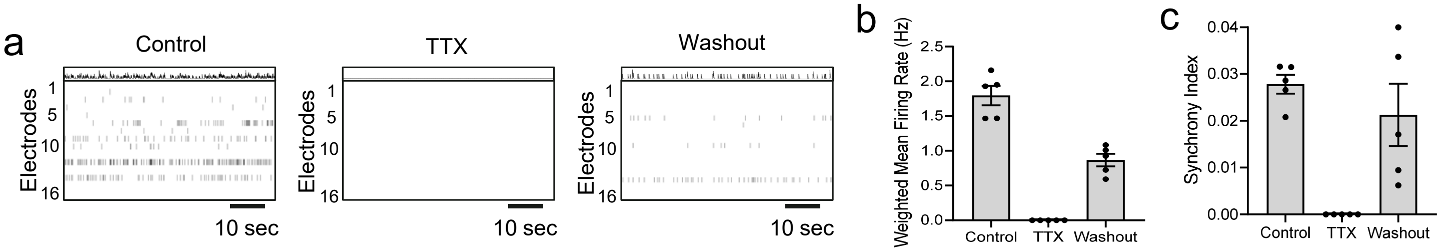

Supplementary Figure 6

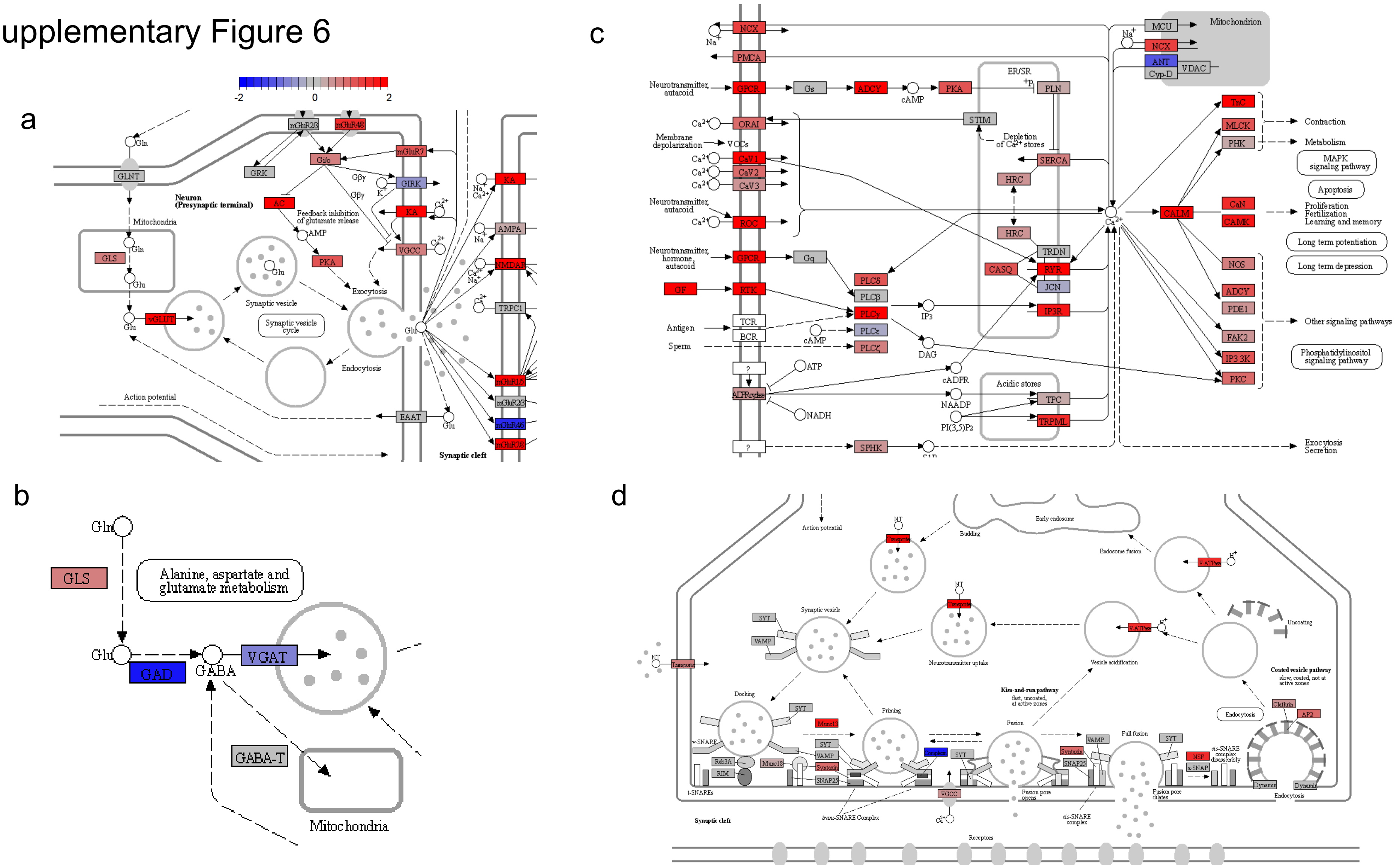
